# Interactive analysis of single-cell data using flexible workflows with SCTK2.0

**DOI:** 10.1101/2022.07.13.499900

**Authors:** Yichen Wang, Irzam Sarfraz, Rui Hong, Yusuke Koga, Vidya Akavoor, Xinyun Cao, Salam Alabdullatif, Nida Pervaiz, Syed Ali Zaib, Zhe Wang, Frederick Jansen, Masanao Yajima, W. Evan Johnson, Joshua D. Campbell

**Author notes:** **Corresponding author:** Joshua D. Campbell, 72 E Concord St, Boston, MA 02118. Authors contributed equally (Yichen Wang, Irzam Sarfraz).

## Abstract

Analysis of single-cell RNA-seq (scRNA-seq) data can reveal novel insights into heterogeneity of complex biological systems. Many tools and workflows have been developed to perform different types of analysis. However, these tools are spread across different packages or programming environments, rely on different underlying data structures, and can only be utilized by people with knowledge of programming languages. In the Single Cell Toolkit 2.0 (SCTK2.0), we have integrated a variety of popular tools and workflows to perform various aspects of scRNA-seq analysis. All tools and workflows can be run in the R console or using an intuitive graphical user interface built with R/Shiny. HTML reports generated with Rmarkdown can be used to document and recapitulate individual steps or entire analysis workflows. We show that the toolkit offers more features when compared with existing tools and allows for a seamless analysis of scRNA-seq data for non-computational users.

**Graphical Abstract:** 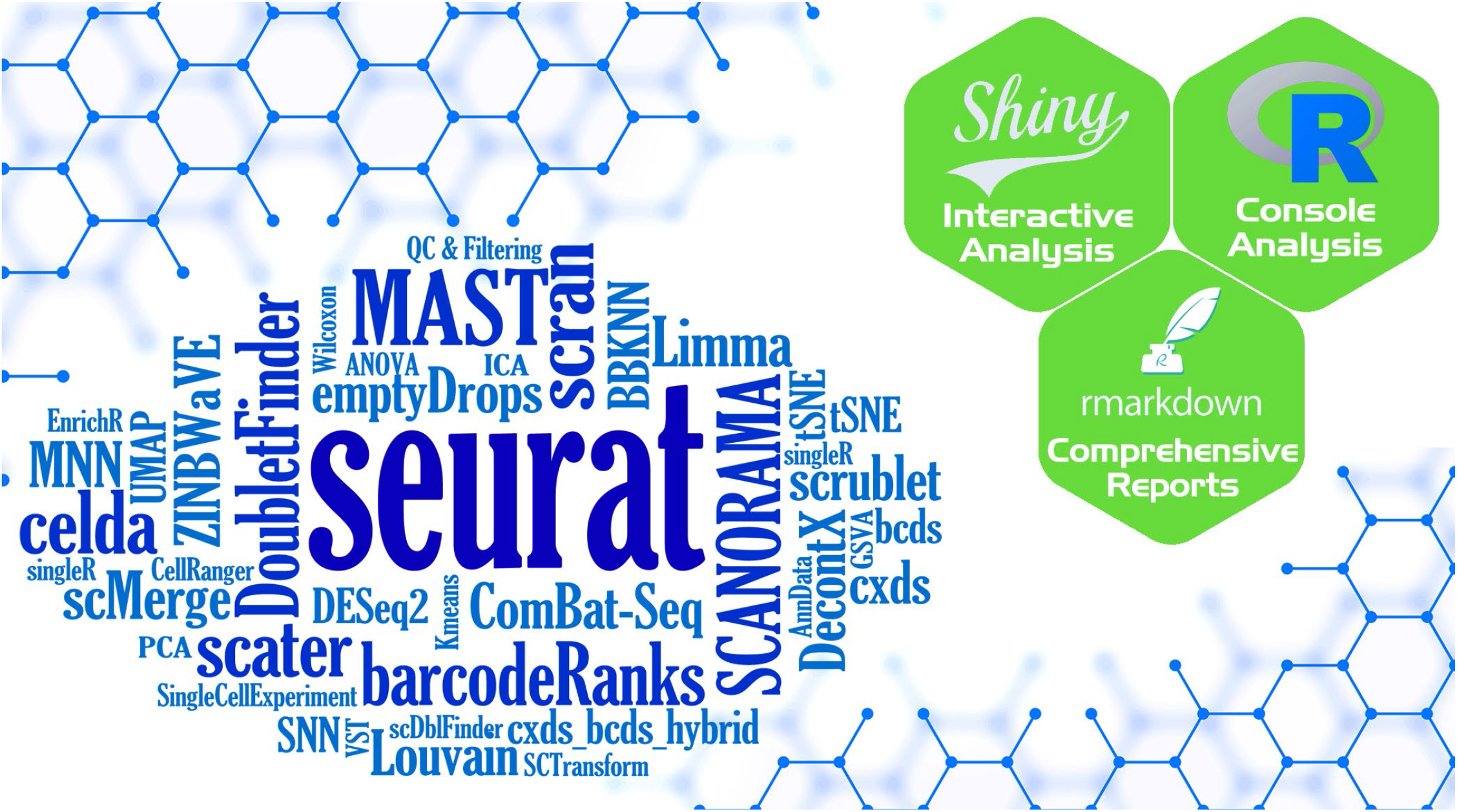

**Highlights:**

- Intuitive graphical user interface for interactive analysis of scRNA-seq data
- Allows non-computational users to analyze scRNA-seq data with end-to-end workflows
- Provides interoperability between tools across different programming environments
- Produces HTML reports for reproducibility and easy sharing of results

## Introduction

Single-cell RNA sequencing (scRNA-seq) is a molecular assay that can quantify of the levels of mRNA transcripts for each gene in individual cells. This approach can be used to generate insights into cellular heterogeneity not previously possible with “bulk” transcriptomic assays [1], [2]. Profiling the transcriptome of individual cells has revealed novel cell subpopulations in normal tissues and cell states associated with the pathogenesis of complex diseases [3]. A large number of tools and software packages are available to perform different steps of scRNA-seq data analysis. However, these tools are spread across different programming environments and rely on different data structures for input of data or output of results. As the interoperability for tools between platforms is lacking, users generally have to choose a single analysis workflow or spend considerable effort manually converting data between environments running different tools and integrating results [4]. Moreover, many researchers without strong computational backgrounds are generating scRNA-seq data but do not have necessary training for analysis and interpretation.

Currently, there are limited options for frameworks that allows for interoperability of tools across environments and contains a graphical user interface (GUI) for non-computational users to perform flexible end-to-end analysis [5][6][7][8]. While some web applications are available for the analysis of scRNA-seq data, there are no online tools that can import data from a variety of formats, perform comprehensive quality control and filtering, run flexible clustering and trajectory workflows, and apply a series of downstream analysis and visualization tools within an interactive interface amiable to users without a strong programming background. To address this need, we developed the Single Cell Toolkit 2.0 (SCTK2.0) which is implemented in the R/Bioconductor package *singleCellTK* and available online at sctk.bu.edu. SCTK2.0 connects our previous R package for quality control of scRNA-seq data [9] with a variety of tools for analysis, integration, and visualization including interoperability with Seurat and many Bioconductor packages. All of the end-to-end analysis workflows are accessible using a “point-and-click” GUI to enable users without programming skills to analyze their own data. When compared to existing tools, the SCTK2.0 framework offers more options for data importing, clustering and trajectory analysis, interactive visualization, and generation of HTML reports for reproducibility.

## Results

### Overview of the general framework

singleCellTK (SCTK) is an R package that provides a uniform interface to popular scRNA-seq tools and workflows for quality control, clustering or trajectory analysis, and visualization. SCTK gives users the opportunity to seamlessly run different tools from different packages and environments during different stages of the analysis. Tools can be run by computational users in the R [10] console, by non-computational users with an interactive graphical user interface (GUI) developed in R/Shiny [11], or with HTML reports generated with Rmarkdown. SCTK utilizes multiple Bioconductor Experiment objects such as the *SingleCellExperiment* (SCE) as the primary data container for storing expression matrices, reduced dimensional representations, cell and feature annotations, and other tool outputs [12][13].

### Flexible and comprehensive workflows for scRNA-seq analysis

The major steps of the SCTK workflow can be divided into three major components: 1) importing, quality control, and filtering, 2) normalization, dimensionality reduction, and clustering, and 3) various downstream analyses and visualizations for exploring biological patterns of the cell clusters (**Fig. 1**). For the first component, we have included the ability to import data from 11 different preprocessing tools or file formats. SCTK generates standard QC metrics such as the total number of counts, features detected per cell, or mitochondrial percentage using the scater package [14]. Doublet detection can be performed with 4 different tools and ambient RNA quantification and removal can be performed with DecontX [15] or SoupX [16]. For filtering, users can choose to exclude cells or genes based on one or a combination of QC metrics produced by the various QC tools.

**Figure 1.**
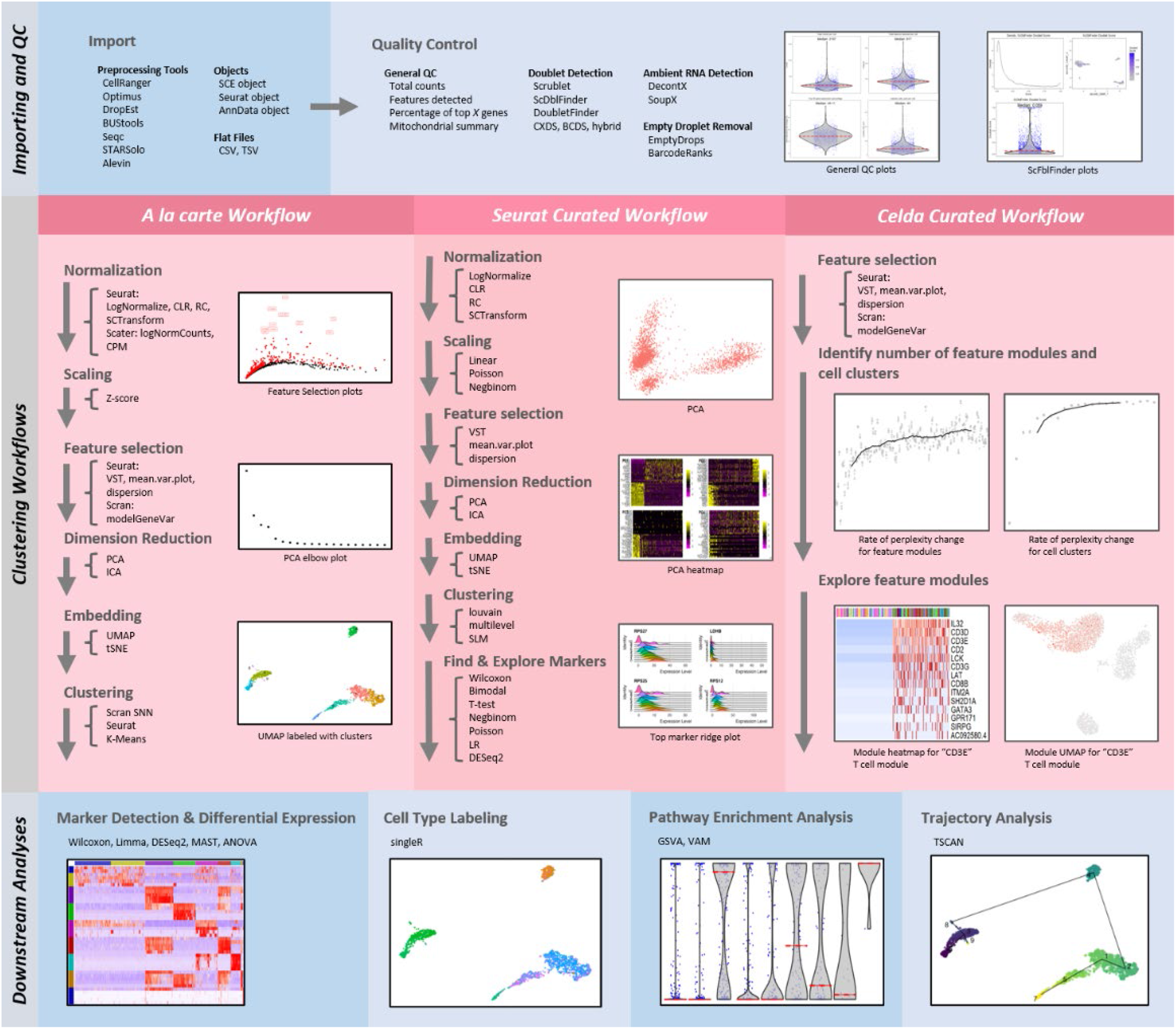
Overview of analysis workflows available in SCTK2.0. Analysis of scRNA-seq data can be divided into three major parts: Importing and Quality Control (QC), Clustering Workflows, and Downstream Analysis. For Importing & QC (top), SCTK2.0 can import data from many different upstream preprocessing tools and formats. A variety of metrics for general QC, empty drop detection, doublet detection, and ambient RNA quantification can be calculated and displayed for each sample. For Clustering Workflows (middle), SCTK2.0 provides an “*a la carte*” workflow which allows users to pick and choose different tools at each step of the workflow as well as curated workflows from the Seurat and Celda packages. For Downstream Analysis (bottom), SCTK2.0 provides access to additional tools and analyses for differential expression, cell type labeling, pathway analysis, and trajectory analysis. Overall, the toolkit provides a wide variety of methods for each part of the analysis workflow.

The major steps for the clustering workflows include normalization, selection of highly variable genes (HVGs), dimensionality reduction such as PCA, clustering, and 2-D embedding such as UMAP (**Fig. 1**). Users also have the option of performing batch correction or integration after normalization with 9 tools. SCTK2.0 provides an “*a la carte”* workflow which allows users to pick and choose different tools at each step or several curated workflows which only allow for specific tools or functions predefined by other packages. Current curated workflows in the Shiny GUI include those from the Seurat [17][18][19][20] and Celda [21] packages.

Downstream analyses after clustering include finding markers for cell clusters using differential expression (DE), DE analysis between user-specified conditions, automated cell type labeling with SingleR [22], pathway enrichment analysis with GSVA [23], VAM [24], or Enrichr [25][26], and trajectory analysis with TSCAN [27]. DE analysis can be performed with the Wilcoxon rank-sum test, MAST [28], Limma [29], ANOVA, or DESeq2 [30] and visualized with heatmaps or volcano plots. The expression of individual genes can be displayed on 2-D embeddings, violin plots, or box plots. Finally, results from SCTK can be exported as flat text files (e.g. mtx, txt, csv), Seurat object, or an AnnData [31][32] object to allow for further analysis and integration with other tools.

### Interactive analysis with the SCTK2.0 GUI

Users without a strong programming background can analyze scRNA-seq data with the interactive GUI built with Shiny and available at sctk.bu.edu (**Fig. 2**). The major steps in the analysis are accessible via the menus in the top navigation bar. Within each major section, parameters to run tools can be selected in the left panel and results are displayed in the right panel. Many plots can be customized with additional options such as the choice of the embedding in a scatter plot or choosing to color the points by a particular metric or label. SCTK has a general visualization tab called the “Cell Viewer” which supports functionality to generate and visualize custom scatter plots, bar plots, and violin plots for user-selected genes or gene sets. Additionally, a generic heatmap plotting tab can be used to visualize the expression levels of multiple features from an expression matrix along with a variety of cell or feature annotations. The majority of plots are made interactive with the *plotly* [33] package and can be highlighted, cropped, zoomed, and saved in various formats.

**Figure 2.**
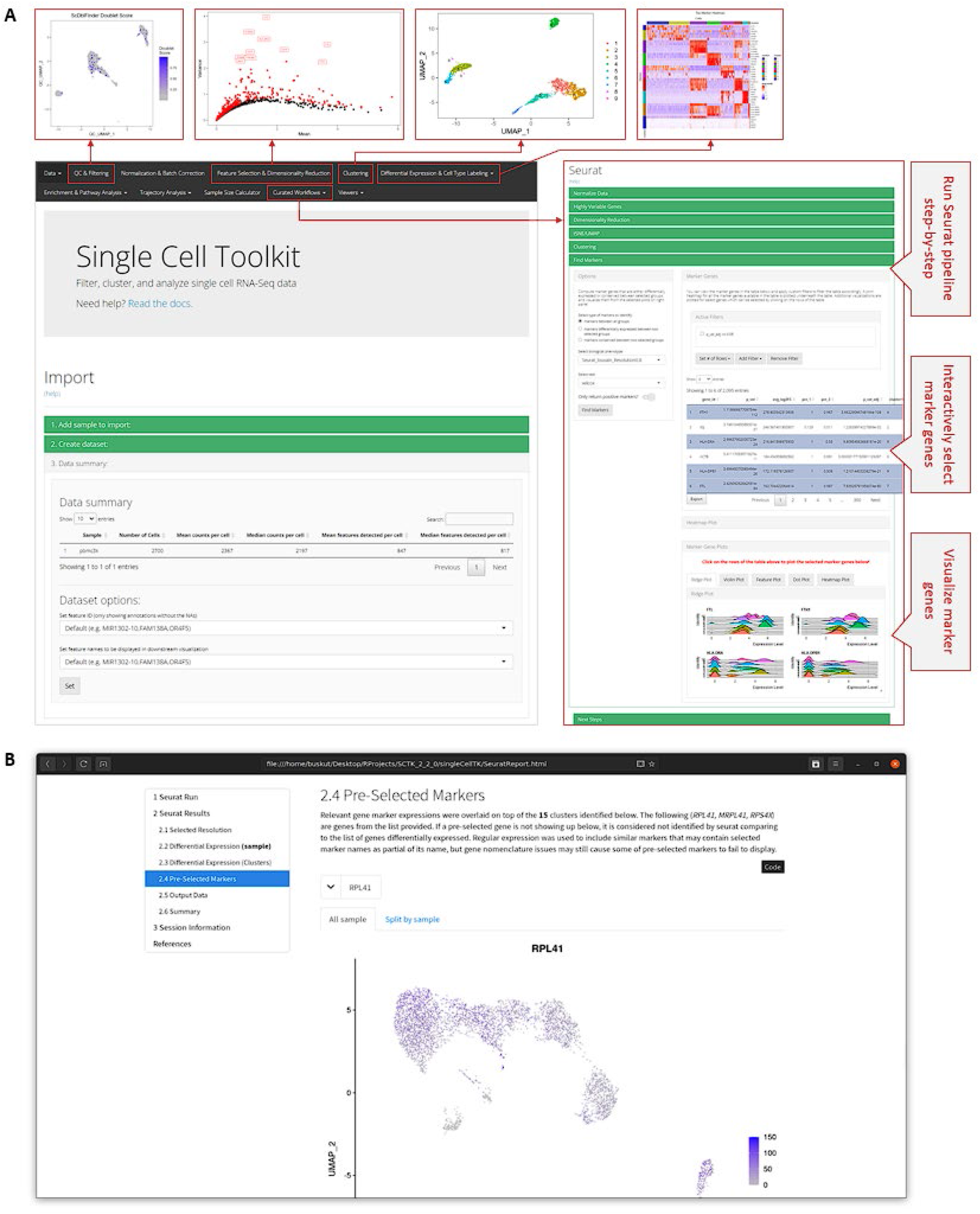
Interactive analysis of single-cell RNA-seq data with a Graphical User Interface (GUI) and HTML reports. SCTK2.0 allows non-computational users to analyze scRNA-seq data using an interactive GUI built with R/Shiny which can be hosted on a web server. **(A)** The menu bar allows the users to navigate through the main sections including data importing, quality control, the “à la carte” clustering workflow, and downstream analysis. The curated workflows for Seurat and Celda can be selected and allow the users to follow through a series of steps using vertical tabs. For example, the Seurat curated workflow is shown and includes steps for normalization, feature selection, dimensionality reduction, clustering, 2-D embedding, and finding markers. **(B)** SCTK2.0 also provides the ability to generate HTML reports for several individual analyses or entire workflows to enable reproducibility and facilitate sharing of results. An HTML report for clustering of PMBC data with Seurat is shown. Different steps that were run in the workflow can be selected with the navigation menu on the left of the report. A description of the step or tool, the chosen parameters, and the resulting plots are shown on the right side of the report.

### Reproducible and sharable analysis with HTML reports

SCTK2.0 can generate HTML reports using Rmarkdown for quality control tools, differential expression (DE) results, differential abundance (DA) results, and for the curated workflows. These reporting tools can be used to plot and share a previously run analysis or start a new analysis workflow *de novo* with user-specified parameters. The output of these functions is a comprehensive HTML report that describes the input data, run parameters, and results with the standard visualizations. These reports provide reproducibility and offer a quick and easy way to explore and share the results of an individual analysis or whole workflow. For example, the DE report renders an HTML document that highlights the top differentially expressed genes via a scrollable table and common visualizations such as a heatmap and volcano plot (**Item S1**).

### Benchmarking

We benchmarked the ability of the SCTK to analyze four datasets of different sizes. Two datasets of peripheral blood mononuclear cells (PBMC) were obtained from 10X Genomics that contained 5,419 cells (*pbmc6k*) and 68,579 cells (*pbmc68k*). Two more datasets of immune cells were obtained from the “1M Immune Cells’’ project from Human Cell Atlas that contained 100k cells (*immune100k)* and 300k cells (*immune300k)*. The workflow consisted of steps of importing data from sparse matrix files, generating QC metrics, filtering, normalization, variable feature selection, dimension reduction, 2D embedding, clustering and marker detection. We recorded the RAM usage for the SCE object after each step (**Fig. S1A**) as well as the peak RAM allocation that was used during each step (**Fig. S1B**). The largest RAM usage for the SCE object was 6.23 GB and occurred after the marker detection step for the largest dataset. The largest peak RAM usage was 16.65 GB and occurred during the importing step of the largest dataset (16.65 GB). These results demonstrate that the SCTK GUI deployed on a server with typical memory availability (e.g. 64GB) can be used to analyze many standard single-cell datasets for several users at a time.

### Comparison to other tools with GUI for scRNA-seq analysis

Some other tools and packages are available that provide graphical user interface to scRNA-seq data analysis. We compared the availability of supported methods between SCTK and Pegasus [5], ASAP [6][7], and BingleSeq [8] (**Table S1**). Generally, SCTK supports more methods and options for the various stages of a typical scRNA-seq analysis. Particularly, SCTK has more options for importing from different data sources and supports more quality control algorithms. Similar to SCTK2.0, several methods and workflows are available in Pegasus. However, the GUI in Pegasus is only available via Jupiter Notebooks in the Terra cloud platform and non-computational users need to have access to a cloud account and a Terra workspace before they can fully utilize this tool. Options for ASAP that are not in SCTK include for voom and DESeq2 for normalization, M3Drop for variable feature detection, and Seurat leiden, hierarchical and SC3 methods for clustering. Lastly, BingleSeq has Monocle for trajectory analysis and dot plots for visualization. With respect to trajectory analyses, SCTK uses TSCAN while Pegasus supports diffusion maps and BingleSeq includes Monocle.

## Discussion

SCTK2.0 provides an intuitive and easy-to-use GUI that integrates a variety of widely used methods into a single end-to-end workflow. Instead of having to switch between different graphic-based tools or learning a programming language to run a method that utilize specific data structures, users can use the “point-and-click” GUI to access existing analysis methods for scRNA-seq data. Features available in the GUI include the ability to import scRNA-seq data from a variety of formats, import and edit annotations for genes and cells, running quality-control analysis and applying filters, applying methods for normalization, dimensionality reduction, clustering, differential expression, pathway analysis, trajectory analysis and interactive visualization. The ability to easily generate comprehensive HTML reports enables quick sharing between collaborators and reproducibility of results. In the future, the *singleCellTK* package will be updated to utilize the *MultiAssayExperiment* and *ExperimentSubset* packages to store and manipulate both multi-modal data and subsets of existing datasets with the same object and from the same interactive interface. Overall, these features make SCTK2.0 a convenient toolkit for the analysis of scRNA-seq data regardless of their programming background.

## STAR★Methods

### Comprehensive Importing

SCTK enables importing data from the following pre-processing tools: CellRanger [34], Optimus, DropEst [35], BUStools [36][37], Seqc [38], STARSolo [39][40] and Alevin [41][42]. In all cases, SCTK parses the standard output directory structure from the pre-processing tools and automatically identifies the count files to import. These functions also support importing of count matrices stored in the plain text files (e.g. MTX, CSV, and TSV formats), SingleCellExperiment (SCE) object saved in RDS file, AnnData object saved in an h5ad file. The Shiny GUI allows users to specify the location of files for multiple samples on their local device. The data for these samples is uploaded and combined into a single SCE object to use across analyses.

### Quality Control and Filtering

Performing comprehensive quality control (QC) is necessary to remove poor quality cells for downstream analysis of single-cell RNA sequencing (scRNA-seq) data. Within droplet-based scRNA-seq data, droplets containing cells must be differentiated from empty droplets. Therefore, assessment of the data is required, for which various QC algorithms have been developed. In SCTK, we support EmptyDrops [43] and BarcodeRank [44] tools for droplets, and general QC Metrics, Scrublet [45], scDblFinder [46], cxds [47], bcds [47], hybrid of cxds and bcds [47], doubletFinder [48] and decontX [15] for cell. The metrics computed from these algorithms can be visualized on a 2D embedding or violin plot. Based on these metrics, users can filter the cells by selecting an appropriate metric and a cutoff value. The filtered data is stored in a separate SCE object and can be utilized in all subsequent analyses.

### À la carte Analysis Workflow

The à la carte analysis workflow includes the main interface and the functions of the toolkit that let the users select and pick different methods and options for various steps of the analysis workflow including normalization, batch correction or integration, feature selection, dimensionality reduction and 2-D embedding, and clustering.

### Normalization

SCTK offers a convenient way to normalize data for downstream analysis using a number of methods available through the toolkit. Normalization methods available with the toolkit include “LogNormalize”, “CLR”, “RC” and “SCTransform” from Seurat package and “logNormCounts” and “CPM” from scater package. Additional transformation options are available for users including “log”, “log1p”, trimming of data assays and Z-Score scaling.

### Batch Correction and Integration

SCTK provides access to methods for batch correction and integration of samples from R packages including Batchelor (MNN) [49], SVA (ComBat) [50][51], limma [29], scMerge [52], Seurat and ZINBWaVE [53], as well as Python packages including BBKNN [54] and Scanorama [55]. These methods accept various types of input expression matrices (e.g. raw counts or log-normalized counts), and generate either a new corrected expression matrix or a low-dimensional dimensionality reduction of the integrated data.

### Feature Selection

Several methods are available to compute and select the most variable features to use in the downstream analysis. Feature selection methods available with the toolkit include “vst”, “mean.var.plot” and “dispersion” from Seurat package and “modelGeneVar” from Scran [56] package. The top variable genes can be visualized through the toolkit in a scatter plot of the genes or features using the mean-to-variance or mean-to-dispersion plot depending upon the algorithm used.

### Dimensionality Reduction and 2D embedding

The toolkit provides access to both PCA (Principal Component Analysis) and ICA (Independent Component Analysis) algorithms from multiple packages for reducing the expression matrices into reduced dimensions. PCA is implemented from both scater and Seurat packages, while implementation of ICA is only available from Seurat. Reduced dimensions computed from these methods can be visualized through various plots including component plot, elbow plot, jackstraw plot and heatmaps. 2D embedding methods available with the toolkit include “tSNE” and “UMAP” from Seurat package, “tSNE” from Rtsne package and “UMAP” from scater package. The results computed from these methods can also be visualized using a 2D scatter plot.

### Clustering

Graph-based clustering methods available within SCTK include “Walktrap” [57], “Louvain” [58], “infomap” [59], “fastGreedy” [60], “labelProp” [61], from the scran package or “louvain”, “multilevel” [62], or “SLM” [63] from the Seurat package. Additionally, K-means methods can be run using “Hartigan-Wong”, “Lloyd”, or “MacQueen” algorithms from the stats package.

### Curated Workflows

SCTK2.0 provides access to both Seurat and Celda analysis workflows through a streamlined and guided interface. Seurat is a widely used R package that implements various methods for processing and clustering of scRNA-seq data. Celda is a R package that performs co-clustering of genes into modules and cells into subpopulations. In the SCTK GUI, all the steps of the Seurat and Celda workflows can be run in a “step-by-step” fashion with the “vertical blinds” layout. These curated workflows allow new or beginner users to quickly run an exploratory analysis of single-cell data without having to try too many combinations of parameters or tools.

### Differential Expression & Marker Selection

The toolkit offers differential expression in a group-vs-group way using one of the five implemented methods including Wilcoxon rank-sum test, MAST, Limma, DESeq2 or ANOVA. Alternatively, users can also use the differential expression methods in a “Find Marker” analysis to identify the top marker genes for each group of cells against all the other cells. The results for both approaches can be viewed through tables that display the top differentially expressed genes or marker genes along with the metrics computed by the selected method.

### Cell Type Labeling

Cell type labeling from a reference can be performed with the SingleR package. SingleR works by comparing the expression profile of each single cell to an annotated reference dataset and labels each cell with a cell type of the highest likelihood. SingleR can also label clusters of cells instead of individual cells. The cell type assignments of clusters or individual cells can be visualized on a 2D embedding in the same fashion as labels from *de novo* clustering algorithms.

### Pathway Analysis

Custom gene sets can be imported by the user or automatically downloaded from the MsigDB [64] database. Methods for scoring the levels of a gene set in each individual cell include Variance-Adjusted Mahalanobis (VAM) and Gene Set Variation Analysis (GSVA). The scores for gene sets can be used in a DE analysis to compare different cell annotations such as cell type or experimental condition. The distribution of gene set scores can be visualized using violin plots. EnrichR can be used to determine if sets of genes are enriched for biological pathways in curated databases such as KEGG [65], GO [66], and MsigDB.

### Trajectory Analysis

Cell trajectory can be constructed by building a cluster based minimum spanning tree (MST) and estimating pseudotime on the paths, with the TSCAN package. Based on the trajectory, SCTK also provides TSCAN methods to test features that are differentially expressed on a path or between paths. The pseudotime value or the expression of DE features can be visualized on a 2D embedding with the MST projected and overlaid on it.

### Benchmarking

The pbmc6k and pbmc68k datasets were obtained using the importExampleData() function which utilized the TENxPBMCData package (version 1.12.0) and ExperimentHub package (version 2.2.1) to retrieve the data. The immune100k and immune300k dataset was retrieved and downsampled from the Human Cell Atlas Portal. All datasets were exported to MTX format. The workflow that was benchmarked included steps for 1) importing the data from an MTX file using the importFromFiles() function, 2) calculation of general quality control metrics using the runPerCellQC() function, 3) normalization using the runNormalization() with the “logNormCounts” method, 4) calculation of variable features using the runFeatureSelection() function with the “modelGeneVar” method, 5) dimensionality reduction using the runDimReduce() function with the “scaterPCA” method, 6) UMAP embedding using the runDimReduce() function with the “scaterUMAP” method, 6) clustering using the runScranSNN() function with the “Louvain” method, and 7) a differential gene expression analysis using the runDEAnalysis() function with the “wilcox” method. For each of the steps, we used the peakRAM() function from the peakRAM package (version 1.0.2) to record the RAM used by the SCE object after the completion of each step as well as the peak RAM allocation used during each step.

## Supporting information

Supplementary Item 1

## Software and Data Availability

Live application: https://sctk.bu.edu/

Documentation and tutorials: https://www.camplab.net/sctk/

Docker image: https://hub.docker.com/r/campbio/sctk_shiny

Bioconductor package: https://bioconductor.org/packages/singleCellTK/

Source code: https://github.com/compbiomed/singleCellTK

*pbmc6k* data: https://support.10xgenomics.com/single-cell-gene-expression/datasets/1.1.0/pbmc6k

*pbmc68k* data: https://support.10xgenomics.com/single-cell-gene-expression/datasets/1.1.0/fresh_68k_pbmc_donor_a

*immune100k* and *immune300k* data: https://data.humancellatlas.org/explore/projects/cc95ff89-2e68-4a08-a234-480eca21ce79

## Author Contributions

Software, Y.W., I.S., R.H., Y.K., V.A., X.C., S.A., N.P., S.A.Z., Z.W., F.J., M.Y., W.E.J. and J.D.C.; Formal Analysis, Y.W. and I.S.; Writing - Original Draft, Y.W. and I.S.; Writing - Review & Editing, J.D.C., W.E.J., Y.K., R.H. Y.W. and I.S., Funding Acquisition, J.D.C., W.E.J. and M.Y.; Supervision, J.D.C.

## Acknowledgements

This work was funded by the National Library of Medicine (NLM) R01LM013154-01 (J.D.C. and M.Y.), the National Cancer Institute (NCI) Informatics Technology for Cancer Research (ITCR) 1U01 CA220413-01 (W.E.J. and J.D.C.), and 5R01GM127430 (W.E.J.).

## Declaration of Interests

The authors declare no competing interests.

## Figures

**Supplementary Figure 1.**
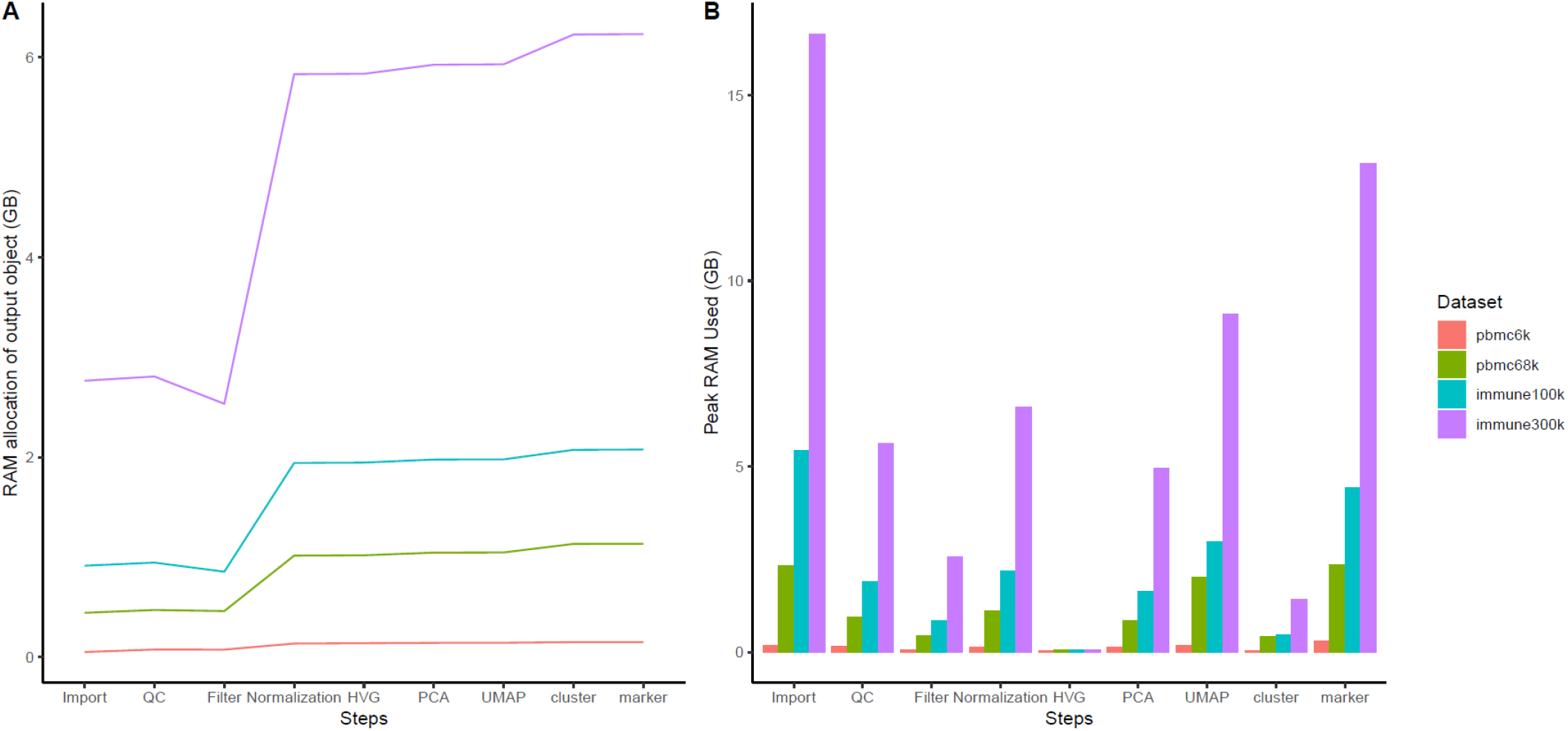
RAM allocation benchmarking for four datasets, *pbmc6k, pbmc68k, immune100k* and *immune300k*, using a Bioconductor based analysis workflow. **A**. The RAM usage for the SCE object after each step is shown for each dataset. **B**. The peak RAM usage during each step is displayed for each dataset.

**Supplementary Table 1.**
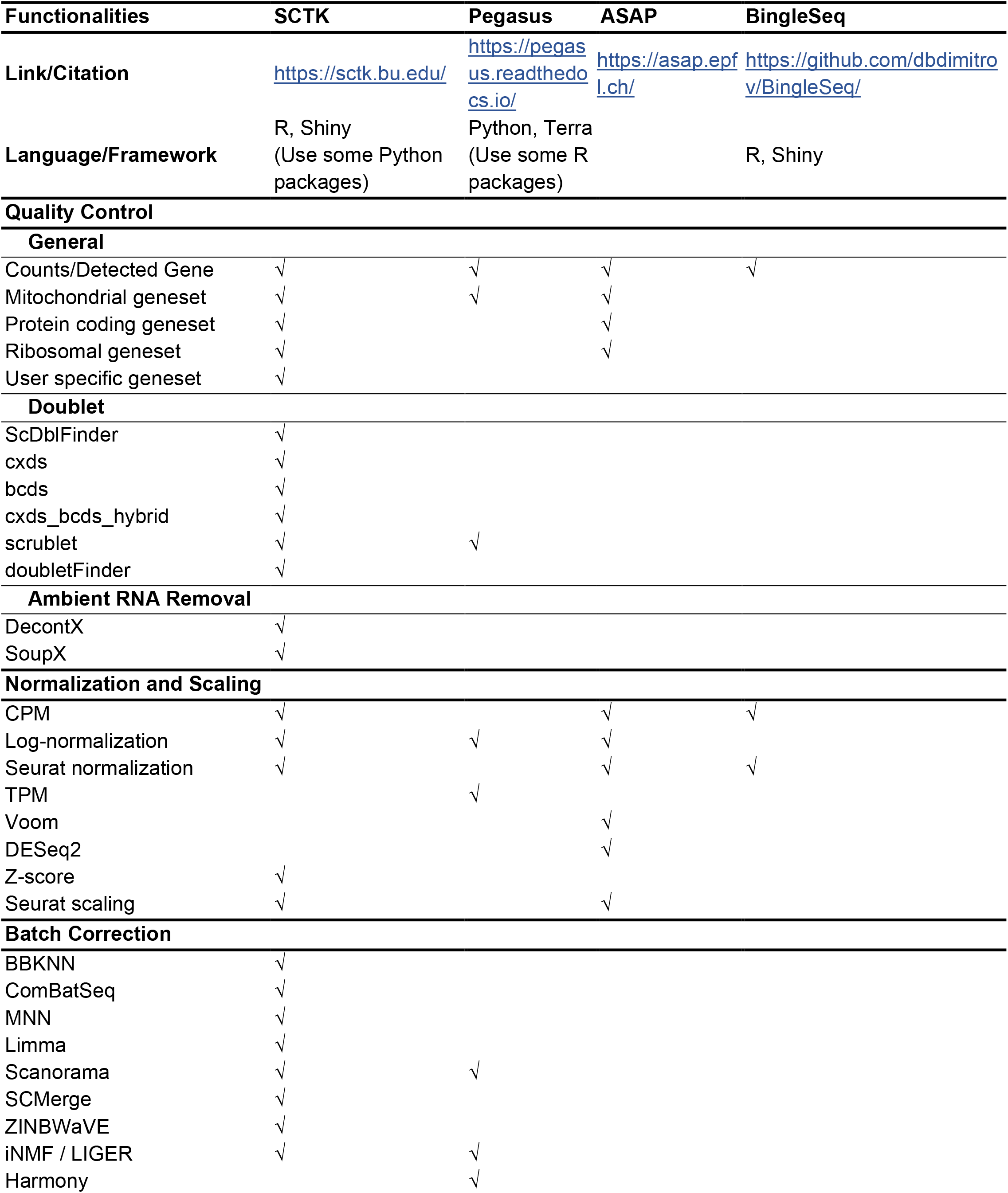
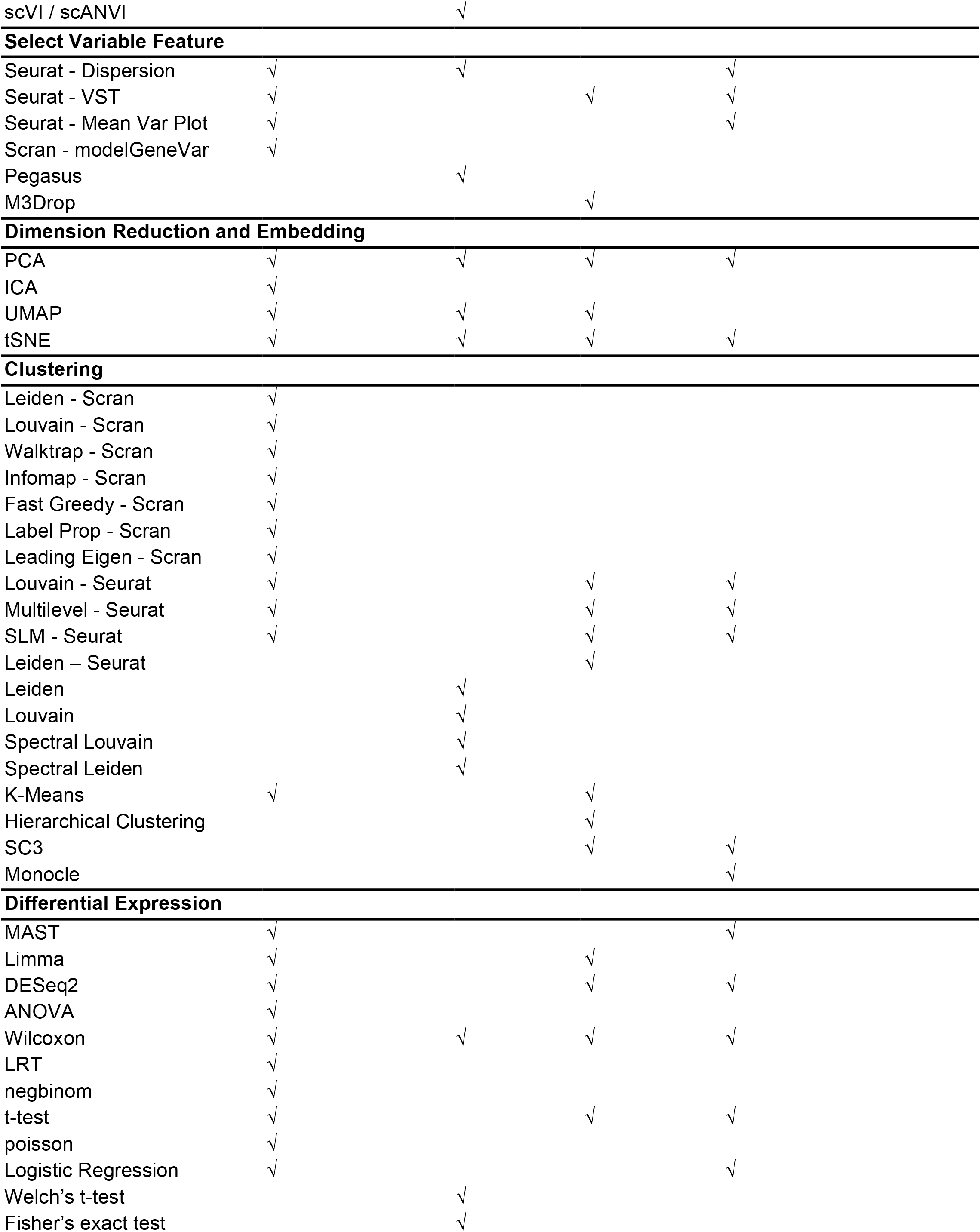
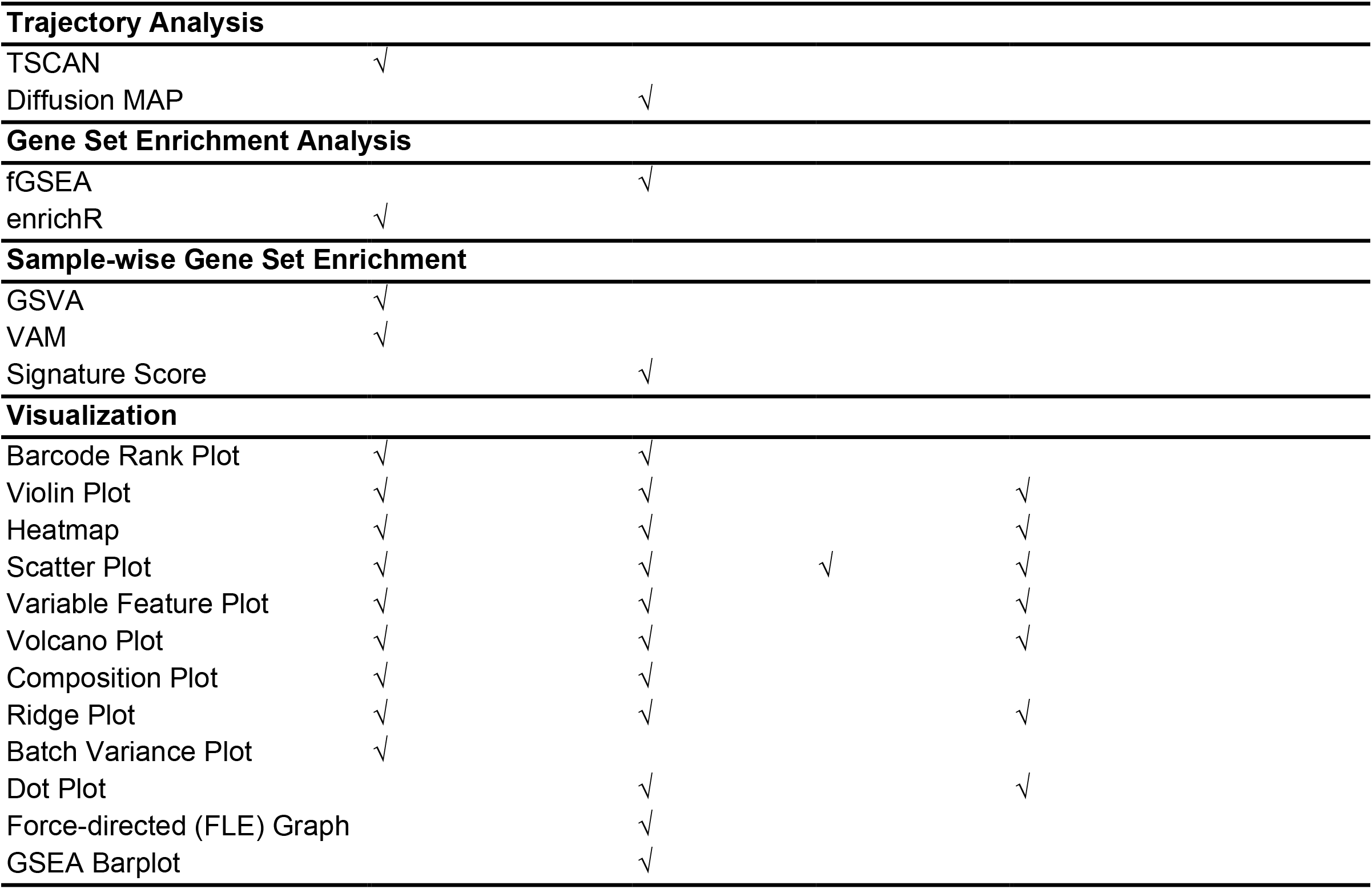

## Notes

### Competing Interest Statement

The authors have declared no competing interest.

## References

[1] Haque, J. Engel, S. A. Teichmann, and T. Lönnberg, “A practical guide to single-cell RNA-sequencing for biomedical research and clinical applications,” Genome Medicine, vol. 9, no. 1, p. 75, Aug. 2017, doi: 10.1186/s13073-017-0467-4.

[2] Hwang, J. H. Lee, and D. Bang, “Single-cell RNA sequencing technologies and bioinformatics pipelines,” Experimental & Molecular Medicine, vol. 50, no. 8, pp. 1–14, Aug. 2018, doi: 10.1038/s12276-018-0071-8.

[3] G. Chen, B. Ning, and T. Shi, “Single-Cell RNA-Seq Technologies and Related Computational Data Analysis,” Frontiers in Genetics, vol. 10, Apr. 2019, doi: 10.3389/fgene.2019.00317.

[4] M. Eisenstein, “Single-cell RNA-seq analysis software providers scramble to offer solutions,” Nature Biotechnology, vol. 38, no. 3, pp. 254–257, Mar. 2020, doi: 10.1038/s41587-020-0449-8.

[5] Li et al., “Cumulus provides cloud-based data analysis for large-scale single-cell and single-nucleus RNA-seq,” Nature Methods, vol. 17, no. 8, pp. 793–798, Aug. 2020, doi: 10.1038/s41592-020-0905-x.

[6] F. P. A. David, M. Litovchenko, B. Deplancke, and V. Gardeux, “ASAP 2020 update: an open, scalable and interactive web-based portal for (single-cell) omics analyses,” Nucleic Acids Research, vol. 48, no. W1, pp. W403–W414, May 2020, doi: 10.1093/nar/gkaa412.

[7] V. Gardeux, F. P. A. David, A. Shajkofci, P. C. Schwalie, and B. Deplancke, “ASAP: a web-based platform for the analysis and interactive visualization of single-cell RNA-seq data,” Bioinformatics, vol. 33, no. 19, pp. 3123–3125, Oct. 2017, doi: 10.1093/bioinformatics/btx337.

[8] Dimitrov and Q. Gu, “BingleSeq: a user-friendly R package for bulk and single-cell RNA-Seq data analysis,” PeerJ, vol. 8, p. e10469, Dec. 2020, doi: 10.7717/peerj.10469.

[9] R. Hong et al., “Comprehensive generation, visualization, and reporting of quality control metrics for single-cell RNA sequencing data,” Nature Communications, vol. 13, no. 1, p. 1688, Dec. 2022, doi: 10.1038/s41467-022-29212-9.

[10] R Core Team, “R: A language and environment for statistical computing.” Vienna, Austria, 2022. [Online]. Available: https://www.R-project.org/

[11] W. Chang et al., “shiny: Web Application Framework for R.” 2021.

[12] R. A. Amezquita et al., “Orchestrating single-cell analysis with Bioconductor,” Nature Methods, vol. 17, no. 2, pp. 137–145, Feb. 2020, doi: 10.1038/s41592-019-0654-x.

[13] I. Sarfraz, M. Asif, and J. D. Campbell, “ExperimentSubset: an R package to manage subsets of Bioconductor Experiment objects,” Bioinformatics, vol. 37, no. 18, pp. 3058–3060, Sep. 2021, doi: 10.1093/BIOINFORMATICS/BTAB179.

[14] J. McCarthy, K. R. Campbell, A. T. L. Lun, and Q. F. Wills, “Scater: Pre-processing, quality control, normalization and visualization of single-cell RNA-seq data in R,” Bioinformatics, vol. 33, no. 8, pp. 1179–1186, Apr. 2017, doi: 10.1093/bioinformatics/btw777.

[15] S. Yang et al., “Decontamination of ambient RNA in single-cell RNA-seq with DecontX,” Genome Biology, vol. 21, no. 1, p. 57, Dec. 2020, doi: 10.1186/s13059-020-1950-6.

[16] M. D. Young and S. Behjati, “SoupX removes ambient RNA contamination from droplet-based single-cell RNA sequencing data,” Gigascience, vol. 9, no. 12, pp. 1–10, Nov. 2020, doi: 10.1093/GIGASCIENCE/GIAA151.

[17] Y. Hao et al., “Integrated analysis of multimodal single-cell data,” Cell, vol. 184, no. 13, pp. 3573–3587.e29, Jun. 2021, doi: 10.1016/J.CELL.2021.04.048.

[18] T. Stuart et al., “Comprehensive Integration of Single-Cell Data Resource Comprehensive Integration of Single-Cell Data,” Cell, vol. 177, 2019, doi: 10.1016/j.cell.2019.05.031.

[19] A. Butler, P. Hoffman, P. Smibert, E. Papalexi, and R. Satija, “Integrating single-cell transcriptomic data across different conditions, technologies, and species,” Nature Biotechnology, vol. 36, no. 5, pp. 411–420, Jun. 2018, doi: 10.1038/nbt.4096.

[20] R. Satija, J. A. Farrell, D. Gennert, A. F. Schier, and A. Regev, “Spatial reconstruction of single-cell gene expression data,” Nature Biotechnology, vol. 33, no. 5, pp. 495–502, May 2015, doi: 10.1038/nbt.3192.

[21] Z. Wang et al., “Celda: A Bayesian model to perform co-clustering of genes into modules and cells into subpopulations using single-cell RNA-seq data,” Biorxiv, Mar. 2021, doi: 10.1101/2020.11.16.373274.

[22] D. Aran et al., “Reference-based analysis of lung single-cell sequencing reveals a transitional profibrotic macrophage,” Nature Immunology 2019 20:2, vol. 20, no. 2, pp. 163–172, Jan. 2019, doi: 10.1038/s41590-018-0276-y.

[23] S. Hänzelmann, R. Castelo, and J. Guinney, “GSVA: gene set variation analysis for microarray and RNA-Seq data,” BMC Bioinformatics, vol. 14, no. 1, p. 7, Jan. 2013, doi: 10.1186/1471-2105-14-7.

[24] H. R. Frost, “Variance-adjusted Mahalanobis (VAM): a fast and accurate method for cell-specific gene set scoring,” Nucleic Acids Research, vol. 48, no. 16, pp. e94–e94, Sep. 2020, doi: 10.1093/NAR/GKAA582.

[25] Y. Chen et al., “Enrichr: Interactive and collaborative HTML5 gene list enrichment analysis tool,” BMC Bioinformatics, vol. 14, no. 1, pp. 1–14, Apr. 2013, doi: 10.1186/1471-2105-14-128.

[26] M. V. Kuleshov et al., “Enrichr: a comprehensive gene set enrichment analysis web server 2016 update,” Nucleic Acids Res, vol. 44, no. W1, pp. W90–W97, Jul. 2016, doi: 10.1093/nar/gkw377.

[27] Z. Ji and H. Ji, “TSCAN: Pseudo-time reconstruction and evaluation in single-cell RNA-seq analysis,” Nucleic Acids Research, vol. 44, no. 13, p. e117, Jul. 2016, doi: 10.1093/nar/gkw430.

[28] Finak et al., “MAST: a flexible statistical framework for assessing transcriptional changes and characterizing heterogeneity in single-cell RNA sequencing data,” Genome Biology, vol. 16, no. 1, p. 278, Dec. 2015, doi: 10.1186/s13059-015-0844-5.

[29] M. E. Ritchie et al., “limma powers differential expression analyses for RNA-sequencing and microarray studies,” Nucleic Acids Research, vol. 43, no. 7, pp. e47–e47, Apr. 2015, doi: 10.1093/nar/gkv007.

[30] M. I. Love, W. Huber, and S. Anders, “Moderated estimation of fold change and dispersion for RNA-seq data with DESeq2,” Genome Biology, vol. 15, no. 12, p. 550, Dec. 2014, doi: 10.1186/s13059-014-0550-8.

[31] I. Virshup, S. Rybakov, F. J. Theis, P. Angerer, and F. A. Wolf, “anndata: Annotated data,” Biorxiv, Dec. 2021, doi: 10.1101/2021.12.16.473007.

[32] F. A. Wolf, P. Angerer, and F. J. Theis, “SCANPY: large-scale single-cell gene expression data analysis,” Genome Biology, vol. 19, no. 1, p. 15, Dec. 2018, doi: 10.1186/s13059-017-1382-0.

[33] C. Sievert, Interactive Web-Based Data Visualization with R, plotly, and shiny. Chapman and Hall/CRC, 2020.

[34] X. Y. Zheng et al., “Massively parallel digital transcriptional profiling of single cells,” Nature Communications, vol. 8, no. 1, p. 14049, Apr. 2017, doi: 10.1038/ncomms14049.

[35] V. Petukhov et al., “dropEst: pipeline for accurate estimation of molecular counts in droplet-based single-cell RNA-seq experiments,” Genome Biology, vol. 19, no. 1, p. 78, Dec. 2018, doi: 10.1186/s13059-018-1449-6.

[36] P. Melsted et al., “Modular, efficient and constant-memory single-cell RNA-seq preprocessing,” Nature Biotechnology, vol. 39, no. 7, pp. 813–818, Jul. 2021, doi: 10.1038/s41587-021-00870-2.

[37] P. Melsted, V. Ntranos, and L. Pachter, “The barcode, UMI, set format and BUStools,” Bioinformatics, vol. 35, no. 21, pp. 4472–4473, Nov. 2019, doi: 10.1093/BIOINFORMATICS/BTZ279.

[38] E. Azizi et al., “Single-Cell Map of Diverse Immune Phenotypes in the Breast Tumor Microenvironment,” Cell, vol. 174, no. 5, pp. 1293–1308.e36, Aug. 2018, doi: 10.1016/j.cell.2018.05.060.

[39] B. Kaminow, D. Yunusov, and A. Dobin, “STARsolo: accurate, fast and versatile mapping/quantification of single-cell and single-nucleus RNA-seq data,” bioRxiv, p. 2021.05.05.442755, May 2021, doi: 10.1101/2021.05.05.442755.

[40] A. Dobin et al., “STAR: ultrafast universal RNA-seq aligner,” Bioinformatics, vol. 29, no. 1, pp. 15–21, Jan. 2013, doi: 10.1093/BIOINFORMATICS/BTS635.

[41] Srivastava, L. Malik, H. Sarkar, and R. Patro, “A Bayesian framework for inter-cellular information sharing improves dscRNA-seq quantification,” Bioinformatics, vol. 36, no. Supplement_1, pp. i292–i299, Jul. 2020, doi: 10.1093/bioinformatics/btaa450.

[42] Srivastava, L. Malik, T. Smith, I. Sudbery, and R. Patro, “Alevin efficiently estimates accurate gene abundances from dscRNA-seq data,” Genome Biology, vol. 20, no. 1, p. 65, Dec. 2019, doi: 10.1186/s13059-019-1670-y.

[43] T. L. Lun, S. Riesenfeld, T. Andrews, T. P. Dao, T. Gomes, and J. C. Marioni, “EmptyDrops: distinguishing cells from empty droplets in droplet-based single-cell RNA sequencing data,” Genome Biology, vol. 20, no. 1, p. 63, Dec. 2019, doi: 10.1186/s13059-019-1662-y.

[44] A. Griffiths, A. C. Richard, K. Bach, A. T. L. Lun, and J. C. Marioni, “Detection and removal of barcode swapping in single-cell RNA-seq data,” Nature Communications, vol. 9, no. 1, p. 2667, Dec. 2018, doi: 10.1038/s41467-018-05083-x.

[45] S. L. Wolock, R. Lopez, and A. M. Klein, “Scrublet: Computational Identification of Cell Doublets in Single-Cell Transcriptomic Data,” Cell Systems, vol. 8, no. 4, pp. 281–291.e9, Apr. 2019, doi: 10.1016/J.CELS.2018.11.005.

[46] P.-L. Germain, A. Lun, C. Garcia Meixide, W. Macnair, and M. D. Robinson, “Doublet identification in single-cell sequencing data using scDblFinder,” F1000Res, vol. 10, p. 979, May 2022, doi: 10.12688/f1000research.73600.2.

[47] A. S. Bais and D. Kostka, “scds: computational annotation of doublets in single-cell RNA sequencing data,” Bioinformatics, vol. 36, no. 4, pp. 1150–1158, Feb. 2020, doi: 10.1093/BIOINFORMATICS/BTZ698.

[48] C. S. McGinnis, L. M. Murrow, and Z. J. Gartner, “DoubletFinder: Doublet Detection in Single-Cell RNA Sequencing Data Using Artificial Nearest Neighbors,” Cell Systems, vol. 8, no. 4, pp. 329–337.e4, Apr. 2019, doi: 10.1016/J.CELS.2019.03.003.

[49] L. Haghverdi, A. T. L. Lun, M. D. Morgan, and J. C. Marioni, “Batch effects in single-cell RNA-sequencing data are corrected by matching mutual nearest neighbors,” Nature Biotechnology, vol. 36, no. 5, pp. 421–427, May 2018, doi: 10.1038/nbt.4091.

[50] T. Leek, W. E. Johnson, H. S. Parker, A. E. Jaffe, and J. D. Storey, “The sva package for removing batch effects and other unwanted variation in high-throughput experiments,” Bioinformatics, vol. 28, no. 6, p. 882, Mar. 2012, doi: 10.1093/BIOINFORMATICS/BTS034.

[51] W. E. Johnson, C. Li, and A. Rabinovic, “Adjusting batch effects in microarray expression data using empirical Bayes methods,” Biostatistics, vol. 8, no. 1, pp. 118–127, Jan. 2007, doi: 10.1093/biostatistics/kxj037.

[52] Y. Lin et al., “scMerge leverages factor analysis, stable expression, and pseudoreplication to merge multiple single-cell RNA-seq datasets,” Proceedings of the National Academy of Sciences, vol. 116, no. 20, pp. 9775–9784, May 2019, doi: 10.1073/pnas.1820006116.

[53] D. Risso, F. Perraudeau, S. Gribkova, S. Dudoit, and J.-P. Vert, “A general and flexible method for signal extraction from single-cell RNA-seq data,” Nature Communications, vol. 9, no. 1, p. 284, Dec. 2018, doi: 10.1038/s41467-017-02554-5.

[54] Polanski, M. D. Young, Z. Miao, K. B. Meyer, S. A. Teichmann, and J. E. Park, “BBKNN: fast batch alignment of single cell transcriptomes,” Bioinformatics, vol. 36, no. 3, pp. 964–965, Feb. 2020, doi: 10.1093/BIOINFORMATICS/BTZ625.

[55] Hie, B. Bryson, and B. Berger, “Efficient integration of heterogeneous single-cell transcriptomes using Scanorama,” Nature Biotechnology, vol. 37, no. 6, pp. 685–691, Jun. 2019, doi: 10.1038/s41587-019-0113-3.

[56] A. T. L. Lun, D. J. McCarthy, and J. C. Marioni, “A step-by-step workflow for low-level analysis of single-cell RNA-seq data with Bioconductor,” F1000Res, vol. 5, p. 2122, Oct. 2016, doi: 10.12688/f1000research.9501.2.

[57] P. Pons and M. Latapy, “Computing Communities in Large Networks Using Random Walks,” in Computer and Information Sciences - ISCIS 2005, vol. 3733, Yolum, T. Güngör, F. Gürgen, and C. Özturan, Eds. Berlin: Springer, 2005, pp. 284–293. doi: 10.1007/11569596_31.

[58] V. D. Blondel, J. L. Guillaume, R. Lambiotte, and E. Lefebvre, “Fast unfolding of communities in large networks,” Journal of Statistical Mechanics: Theory and Experiment, vol. 2008, no. 10, p. P10008, Oct. 2008, doi: 10.1088/1742-5468/2008/10/P10008.

[59] Rosvall, D. Axelsson, and C. T. Bergstrom, “The map equation,” The European Physical Journal Special Topics, vol. 178, no. 1, pp. 13–23, Nov. 2009, doi: 10.1140/epjst/e2010-01179-1.

[60] A. Clauset, M. E. J. Newman, and C. Moore, “Finding community structure in very large networks,” Physical Review E, vol. 70, no. 6, p. 066111, Dec. 2004, doi: 10.1103/PhysRevE.70.066111.

[61] X. Zhu and Z. Ghahramani, “Learning from Labeled and Unlabeled Data with Label Propagation,” Carnegie Mellon University, Pittsburgh, 2002.

[62] R. Rotta and A. Noack, “Multilevel local search algorithms for modularity clustering,” ACM Journal of Experimental Algorithmics, vol. 16, May 2011, doi: 10.1145/1963190.1970376.

[63] Waltman and N. J. van Eck, “A smart local moving algorithm for large-scale modularity-based community detection,” The European Physical Journal B, vol. 86, no. 11, p. 471, Nov. 2013, doi: 10.1140/epjb/e2013-40829-0.

[64] A. Liberzon, C. Birger, H. Thorvaldsdóttir, M. Ghandi, J. P. Mesirov, and P. Tamayo, “The Molecular Signatures Database Hallmark Gene Set Collection,” Cell Systems, vol. 1, no. 6, pp. 417–425, Dec. 2015, doi: 10.1016/j.cels.2015.12.004.

[65] Kanehisa and S. Goto, “KEGG: Kyoto Encyclopedia of Genes and Genomes,” Nucleic Acids Research, vol. 28, no. 1, pp. 27–30, Jan. 2000, doi: 10.1093/nar/28.1.27.

[66] Gene Ontology Consortium, “The Gene Ontology (GO) database and informatics resource,” Nucleic Acids Research, vol. 32, no. 90001, pp. D258–D261, Jan. 2004, doi: 10.1093/nar/gkh036.

