## Supplementary Item 1 for "Interactive analysis of single-cell data using flexible workflows with SCTK2.0": Supplementary Item 1.html

Differential Expression Analysis


Code 

- Show All Code
- Hide All Code

### Differential Expression Analysis

###### July 11, 2022

### Analysis: T cell subpopulations

The differential expression was performed by comparing the feature
expression of cells belonging to condition “**cluster1**”
against the cells belonging to condition “**cluster2**”.
The relationship between the two selected group is could be indicated
via the embedding plot below. The feature expression matrix used was
“**logcounts**”.

In the condition “**cluster1**”, **560**
cells were included, while **538** cells were included for
condition **cluster2**.

The method used for performing the differential expression analysis
was “**wilcox**”. For the more information of the method,
please refer to the
help page.

```
sce$deClass <- "Not involved"
sce$deClass[metas$select$ix1] <- cond1
sce$deClass[metas$select$ix2] <- cond2
twoColors <- discreteColorPalette(2, palette = "ggplot")
colorValue <- c("grey", twoColors[1], twoColors[2])
names(colorValue) <- c("Not involved", cond1, cond2)
plotSCEDimReduceColData(sce, colorBy = "deClass", reducedDimName = useReducedDim, labelClusters = FALSE) + scale_color_manual(values = colorValue)
```

#### Result Table

The following table presents the statistic metrics of the
differential expression analysis. The DEGs in the table are pre-filtered
by FDR value less than 0.05.

```
resultTable <- getDEGTopTable(sce, useResult = study, log2fcThreshold = 0, labelBy = featureDisplay)
datatable(resultTable, options = list(pageLength = 10, scollX = 400))
```

> The table above can be reordered by clicking on the column name.

### Visualization

#### Heatmap

The following heatmap will be divided by the conditions and the
regulation. A gene will be determined as up-regulated in condition
**cluster1** if its log2FC value was greater than 0.25 and
FDR is less than 0.05, and down-regulated in condition
**cluster1** if the log2FC value is less than -0.25. The
gene regulation and condition setting will be annotated by default. In
addition, if the condition setting was achieved by using a categorical
annotation in the object, the annotation used will also be labeled to
the column level of the heatmap.

```
plotDEGHeatmap(sce, study, rowLabel = featureDisplay)
```

#### Volcano Plot

The following volcano plot contains dots as all the genes, while the
log2FC value on the X axis and -log10(FDR) value on the Y axis. That’s
to say, the genes farther away from the center of the plot (i.e. at the
left or right) show larger fold-change between the two conditions, while
the genes closer to the top of the plot show higher significance. The
genes with positive log2FC values are up-regulated in the group
**cluster1**, and those with negative log2FC values are
up-regulated in **cluster2**. 10 genes with top fold-change
(absolute value) are labeled with text.

```
plotDEGVolcano(sce, study, featureDisplay = featureDisplay)
```

#### Regression Plot

The following plot shows the linear model regression of the top 36
DEGs, ranked by FDR values.

```
plotDEGRegression(sce, study, labelBy = featureDisplay)
```

#### Violin Plot

The following plot shows the violin plot also for comparing the
expression of the top 36 DEGs between the two conditions.

```
plotDEGViolin(sce, study, labelBy = featureDisplay)
```

### SessionInfo

```
sessionInfo()
```

```
## R version 4.2.0 (2022-04-22 ucrt)
## Platform: x86_64-w64-mingw32/x64 (64-bit)
## Running under: Windows 10 x64 (build 19043)
## 
## Matrix products: default
## 
## locale:
## [1] LC_COLLATE=Chinese (Simplified)_China.utf8  LC_CTYPE=Chinese (Simplified)_China.utf8   
## [3] LC_MONETARY=Chinese (Simplified)_China.utf8 LC_NUMERIC=C                               
## [5] LC_TIME=English_United States.1252         
## 
## attached base packages:
## [1] grid      stats4    stats     graphics  grDevices utils     datasets  methods   base     
## 
## other attached packages:
##  [1] DT_0.23                     patchwork_1.1.1             gridExtra_2.3              
##  [4] scater_1.24.0               scuttle_1.6.2               kableExtra_1.3.4           
##  [7] knitr_1.39                  ggplot2_3.3.6               RColorBrewer_1.1-3         
## [10] cowplot_1.1.1               dplyr_1.0.9                 sp_1.5-0                   
## [13] SeuratObject_4.1.0          Seurat_4.1.1                TENxPBMCData_1.14.0        
## [16] HDF5Array_1.24.1            rhdf5_2.40.0                singleCellTK_2.7.1         
## [19] DelayedArray_0.22.0         Matrix_1.4-1                SingleCellExperiment_1.18.0
## [22] SummarizedExperiment_1.26.1 Biobase_2.56.0              GenomicRanges_1.48.0       
## [25] GenomeInfoDb_1.32.2         IRanges_2.30.0              S4Vectors_0.34.0           
## [28] BiocGenerics_0.42.0         MatrixGenerics_1.8.0        matrixStats_0.62.0         
## 
## loaded via a namespace (and not attached):
##   [1] rappdirs_0.3.3                MCMCprecision_0.4.0           scattermore_0.8              
##   [4] R.methodsS3_1.8.2             tidyr_1.2.0                   bit64_4.0.5                  
##   [7] irlba_2.3.5                   R.utils_2.12.0                data.table_1.14.2            
##  [10] rpart_4.1.16                  KEGGREST_1.36.2               RCurl_1.98-1.7               
##  [13] doParallel_1.0.17             generics_0.1.3                ScaledMatrix_1.4.0           
##  [16] RSQLite_2.2.14                RANN_2.6.1                    combinat_0.0-8               
##  [19] future_1.26.1                 bit_4.0.4                     spatstat.data_2.2-0          
##  [22] webshot_0.5.3                 xml2_1.3.3                    httpuv_1.6.5                 
##  [25] assertthat_0.2.1              viridis_0.6.2                 xfun_0.31                    
##  [28] jquerylib_0.1.4               evaluate_0.15                 promises_1.2.0.1             
##  [31] fansi_1.0.3                   assertive.files_0.0-2         dbplyr_2.2.1                 
##  [34] igraph_1.3.2                  DBI_1.1.3                     htmlwidgets_1.5.4            
##  [37] spatstat.geom_2.4-0           purrr_0.3.4                   ellipsis_0.3.2               
##  [40] crosstalk_1.2.0               deldir_1.0-6                  sparseMatrixStats_1.8.0      
##  [43] vctrs_0.4.1                   ROCR_1.0-11                   abind_1.4-5                  
##  [46] cachem_1.0.6                  RcppEigen_0.3.3.9.2           withr_2.5.0                  
##  [49] GSVAdata_1.32.0               svMisc_1.2.3                  progressr_0.10.1             
##  [52] sctransform_0.3.3             goftest_1.2-3                 svglite_2.1.0                
##  [55] cluster_2.1.3                 ExperimentHub_2.4.0           lazyeval_0.2.2               
##  [58] crayon_1.5.1                  edgeR_3.38.1                  pkgconfig_2.0.3              
##  [61] labeling_0.4.2                nlme_3.1-157                  vipor_0.4.5                  
##  [64] rlang_1.0.2                   globals_0.15.1                lifecycle_1.0.1              
##  [67] miniUI_0.1.1.1                filelock_1.0.2                BiocFileCache_2.4.0          
##  [70] enrichR_3.0                   rsvd_1.0.5                    AnnotationHub_3.4.0          
##  [73] polyclip_1.10-0               lmtest_0.9-40                 Rhdf5lib_1.18.2              
##  [76] zoo_1.8-10                    beeswarm_0.4.0                GlobalOptions_0.1.2          
##  [79] ggridges_0.5.3                rjson_0.2.21                  png_0.1-7                    
##  [82] viridisLite_0.4.0             bitops_1.0-7                  R.oo_1.25.0                  
##  [85] KernSmooth_2.23-20            rhdf5filters_1.8.0            Biostrings_2.64.0            
##  [88] blob_1.2.3                    DelayedMatrixStats_1.18.0     shape_1.4.6                  
##  [91] stringr_1.4.0                 parallelly_1.32.0             spatstat.random_2.2-0        
##  [94] gridGraphics_0.5-1            beachmat_2.12.0               scales_1.2.0                 
##  [97] memoise_2.0.1                 magrittr_2.0.3                plyr_1.8.7                   
## [100] ica_1.0-3                     zlibbioc_1.42.0               compiler_4.2.0               
## [103] dqrng_0.3.0                   clue_0.3-61                   fitdistrplus_1.1-8           
## [106] cli_3.3.0                     XVector_0.36.0                listenv_0.8.0                
## [109] pbapply_1.5-0                 MASS_7.3-57                   mgcv_1.8-40                  
## [112] tidyselect_1.1.2              MAST_1.22.0                   stringi_1.7.6                
## [115] highr_0.9                     yaml_2.3.5                    assertive.numbers_0.0-2      
## [118] BiocSingular_1.12.0           locfit_1.5-9.5                ggrepel_0.9.1                
## [121] sass_0.4.1                    tools_4.2.0                   future.apply_1.9.0           
## [124] parallel_4.2.0                circlize_0.4.15               rstudioapi_0.13              
## [127] foreach_1.5.2                 celda_1.12.0                  assertive.types_0.0-3        
## [130] farver_2.1.1                  Rtsne_0.16                    DropletUtils_1.16.0          
## [133] digest_0.6.29                 BiocManager_1.30.18           rgeos_0.5-9                  
## [136] shiny_1.7.1                   Rcpp_1.0.8.3                  BiocVersion_3.15.2           
## [139] later_1.3.0                   RcppAnnoy_0.0.19              httr_1.4.3                   
## [142] AnnotationDbi_1.58.0          ComplexHeatmap_2.12.0         assertive.properties_0.0-5   
## [145] colorspace_2.0-3              rvest_1.0.2                   tensor_1.5                   
## [148] reticulate_1.25               splines_4.2.0                 uwot_0.1.11                  
## [151] spatstat.utils_2.3-1          plotly_4.10.0                 systemfonts_1.0.4            
## [154] xtable_1.8-4                  assertive.base_0.0-9          jsonlite_1.8.0               
## [157] R6_2.5.1                      pillar_1.7.0                  htmltools_0.5.2              
## [160] mime_0.12                     glue_1.6.2                    fastmap_1.1.0                
## [163] BiocParallel_1.30.3           BiocNeighbors_1.14.0          interactiveDisplayBase_1.34.0
## [166] codetools_0.2-18              fishpond_2.2.0                utf8_1.2.2                   
## [169] lattice_0.20-45               bslib_0.3.1                   spatstat.sparse_2.1-1        
## [172] tibble_3.1.7                  multipanelfigure_2.1.2        curl_4.3.2                   
## [175] ggbeeswarm_0.6.0              leiden_0.4.2                  gtools_3.9.2.2               
## [178] magick_2.7.3                  survival_3.3-1                limma_3.52.2                 
## [181] rmarkdown_2.14                munsell_0.5.0                 GetoptLong_1.0.5             
## [184] GenomeInfoDbData_1.2.8        iterators_1.0.14              reshape2_1.4.4               
## [187] gtable_0.3.0                  spatstat.core_2.4-4
```
